# Introducing entropy-based metrics for quantifying edge- and macro-shape complexity in leaves and beyond

**DOI:** 10.64898/2026.08.26.747315

**Authors:** Tobias Trauden, Anjaharinony A. N. A. Rakotomalala, Robert R. Junker, Lucy Sauereßig, Kerima Trauden, Merle Muñoz Andres, Rebecca Dannoritzer, Nina Farwig, Stefan Pinkert

## Abstract

Leaf shape is a fundamental trait of plant ecological strategies, influencing biotic interactions and ecosystem functioning. However, established quantitative metrics fail to capture subtle variations and irregularities, require user-based reference points or are challenging to compare among taxa with broadly different leaf shapes. In addition, established metrics typically conflate (aggregate) leaf edge complexity and macro-shape complexity, despite their independent functional significance and genetic foundations. Here, we introduce an entropy-based framework to quantify two new complexity metrics: edge complexity and macro-shape complexity. Based on three case studies, we show that these metrics outperform aggregate metrics in predicting *Quercus robur* chemical traits, provide more intuitive interspecific classifications, and strongly align with human perception. In addition, edge and macro-shape complexity show high complementarity, while aggregate metrics are highly redundant and typically strongly related to leaf area. Emerging as the strongest predictor of leaf chemistry and key visual cue for complexity as perceived by humans, the effects of edge complexity highlight the under-appreciated functional significance of leaf margins. Our framework and the proposed entropy-based complexity metrics thus promise to help unlock the potential of growing digital image archives of leaves, including images from herbaria and fossils, and are technically readily applicable to shapes of algae, bacteria, pollen, and beyond. The accompanying package *ShapeComplexity* enables the broad application of entropy-based metrics, providing a powerful tool to explore how the shape of organisms and biological structures influences ecological strategies, biotic interactions, and ecosystem functioning while tracking spatial and temporal variation.

## INTRODUCTION

Functional traits are fundamental to understanding species’ distributions, local adaptations, and interactions with both abiotic and biotic factors across space and time (Mcgill *et al*. 2006; Violle *et al*. 2007). Plants offer an exceptionally rich source of functional trait data. This wealth of biodiversity data is the result of a long-standing interest in the ecological and evolutionary role of trait variation and a growing use of automated measurement approaches (Kattge *et al*. 2020). Among the most readily available and commonly measured traits are leaf area, circularity, and shape index, which typically serve as proxies for photosynthetic capacity, productivity (Givnish 1979; Tsukaya 2006, 2018), heat load, and water loss (Leigh *et al*. 2017; Nicotra *et al*. 2011; Peppe *et al*. 2011). Together with the symmetry and irregularity of leaves as characterised by fractal dimension (Mandelbrot & Wheeler 1983) (Herrera 2017) shape metrics have attracted renewed interest during the last decade for inferring responses to environmental stress or herbivory, including their spatial and temporal trends (Arseniou & MacFarlane 2021; Li *et al*. 2020). However, established shape metrics primarily emphasize broad, interspecific differences in overall shape, reducing the remarkable diversity of leaf forms to coarse distinctions (Nicotra *et al*. 2011; Royer *et al*. 2005), while subtle but functionally relevant variation often escapes detection (Chitwood & Sinha 2016; Li et al. 2020).

Well recognised for its ecological and evolutionary importance, leaf shape variation is ubiquitous both among and within species (Bhatia *et al*. 2021; Nakayama 2024). Complexity, defined by Lorenz (Lorenz 1993) as “irregularity in space” or the length of the “set of instructions” provides an ordered measure of shape variation that can be readily applied to overall and component patterns. In biological systems, complexity equates to the presence and abundance of sophisticated structures (Mitchell 2011), such as edge features (fine-scale features like entire, toothed, or serrated edges) and macro-shape features (general shape features like round, lanceolate, or lobed forms) of leaves – features that are notably widespread (Givnish 1987; Peppe *et al*. 2011). For instance, up to 80% of all and 64% of the most common tree species in Germany have pronounced lobes, leaflets, or toothed macro-shapes (Holzwarth *et al*. 2020).

The remarkable macros-shape diversity is the primary focus of ecological research and is central to taxonomy and modern image-based species classification systems (Cope *et al*. 2012). However, many broad-leaved species also display multiple leaf shapes (i.e., heterophylly) and nuanced intraspecific variation associated with abiotic drivers and biotic interactions. For example, sun leaves frequently exhibit greater dissection than shade leaves of the same individual, improving boundary-layer conductance and reducing water loss (Leigh *et al*. 2017). More broadly, intraspecific leaf shape variation can serve as an accessible proxy for variation in chemical traits (such as chlorophyll, flavanol, anthocyanin, and nitrogen content) due to their shared dependence on resource allocation and stress physiology (Pérez-Harguindeguy *et al*. 2013). Intraspecific variation in leaf shape also mediates ecological interactions with some herbivorous insects using leaf outlines as visual cues for host recognition (Rausher 1978; Schoonhoven *et al*. 2010), while others are directly affected by shape-dependent defences (Higuchi & Kawakita 2019). Collectively, these examples underscore the potential of shape analysis for elucidating the ecological and functional role of complexity, but they also highlight challenges regarding its quantification.

Although many existing shape metrics are easy to measure, either manually or automatically, they primarily capture coarse differences in length and area and are therefore relatively insensitive to fine-scale edge structure. As a result, and despite their functional importance and largely independent genetic control, edge features remain underrepresented in quantitative analyses of leaf morphology (Chitwood & Sinha 2016; Li *et al*. 2020). Human-based classifications remain the most common means of distinguishing macro-from edge-shape variation, but they rely on nominal categories such as “lobed,” “serrated,” or “entire.” More precise landmark-based methods capture greater detail, but they are labour-intensive and often difficult to reproduce (Kozlov *et al*. 2017). Although these limitations of landmark-based methods can be addressed using Elliptical Fourier Transform (Klein & Svoboda 2017), which does not require user-input, both approaches are mechanistically ambiguous and offer limited translational applicability, thereby constraining comparisons across study systems and among species with substantial intraspecific variation, such as those displaying heterophylly. Indeed, intraspecific variation is increasingly recognised as an integral component of ecological strategies and community dynamics rather than statistical noise (Albert *et al*. 2010; Violle *et al*. 2012). Hence, the reliance on a fixed and limited set of shape categories, landmarks or transformation vectors, rather than universal quantitative metrics, critically hampers our understanding of the mechanisms driving shape variation and its ecological and evolutionary significance (Nakayama 2024).

Entropy-based approaches provide a promising alternative to widely used shape metrics, such as circularity and shape index. Importantly, entropy-based metrics can be applied to separate features of an object tailored to reflect different ecological functions and foundations. On the one hand, they can address both the amplitude and regularity of edge features. For instance, the approximate entropy of a signal curve retrieved for any given linearised structure can measure the degree of disorder or unpredictability in a system (Marcon & Puech 2017; Shannon 1948; Wu *et al*. 2013), similar to the concept of fractal dimension, which is technically restricted to descriptions of entire objects (Berntson & Stoll 1997; Mandelbrot 1967). On the other hand, the low sensitivity of spectral entropy to outliers makes it particularly well suited for capturing macro-shape complexity in terms of the overall frequency distribution of shape variation.

To overcome the methodological and conceptual limitations of existing shape metrics (Table 1), here we introduce a novel analytical framework that quantifies the two main dimensions of complexity. Our approach is based on the approximate entropy of a linearised signal curve of fine-scale edge features and the spectral entropy of the signal curve of leaf macro-shape features. Our methodological framework, implemented in the software package *ShapeComplexity*, is designed to enable rapid and reproducible computation of these new metrics of leaf edge complexity (EC) and leaf macro-shape complexity (MC) as well as a range of conventional metrics that we use to evaluate our method in three case studies. First, we assess the contribution of entropy-based metrics to intraspecific shape variation and their ability to predict chemical leaf traits using over 1100 leaves from eight *Quercus robur* trees –a species that shows extreme heterophylly from round to lobed leaves with entire and toothed edges– in similar environments. Second, we test their ability to differentiate species and genera among 13 deciduous tree species across Sweden –covering toothed, serrated, lobed, round and lacteous species. Third, we evaluate the association between aggregate and entropy-based metrics and leaf shape complexity as assessed by humans, hypothesizing that entropy-based metrics better capture the nuanced irregularities and structural frequencies that underpin intuitive assessments of complexity relative to traditional categorizations of leaf shape.

**Table 1:** Comparative evaluation of commonly used approaches for quantifying shape complexity based on methodological properties. Criteria emphasize scalability, generalizability, and robustness to measurement conditions, including image resolution (summarizing references from the introduction). *User input required* refers to the degree of manual intervention necessary to parameterize the method prior to analysis, such as placement of homologous landmarks or assignment of categorical descriptors. High values indicate substantial manual effort and potential observer dependence. *Extrapolation capacity* (transferability) denotes the capacity of a fitted representation or classification scheme to be applied to novel shapes without redefining the underlying model structure or feature set. Methods with high transferability rely on general geometric properties rather than predefined biological features and therefore support extrapolation across morphologically disparate taxa. *Automation* describes the extent to which the method can be executed algorithmically once input data (e.g., binary outlines) are available. Highly automated methods are generally more scalable and reproducible across large datasets. *Aggregate metric* indicates whether the method reduces shape complexity to a single summary statistic that does not explicitly distinguish among sources of variation, such as global form versus boundary structure. Aggregate metrics are computationally efficient but typically contain limited structural information. Consistent *biological interpretability* reflects the degree to which a method preserves spatial or structural information that can be interpreted in functional, developmental, or evolutionary terms. High mechanistic content implies retention of biologically meaningful geometry rather than compression into a scalar descriptor. *Area-dependence* (per se) describes whether metric values are intrinsically influenced by object size or scale. Metrics with high dependence on area may confound size and complexity unless explicitly normalized. *Quantitative vs. qualitative* distinguishes continuous numerical descriptors from categorical classifications. Quantitative metrics enable statistical modeling and hypothesis testing, whereas qualitative approaches primarily support descriptive or diagnostic assessments. *Irregularity sensitivity* refers to the sensitivity of the method to fine-scale deviations from smooth boundaries, such as serrations, lobes, or hierarchical edge complexity. Methods with high capability can capture variation in boundary roughness across spatial scales.

| Method | User input required | Extrapolation capacity | Automation | Aggregate metric | Biological interpretability | Area-dependence | Quantitative vs. qualitative | Irregularity sensitivity |
| --- | --- | --- | --- | --- | --- | --- | --- | --- |
| Landmark-based morphometrics | High | Low | Low–Moderate | No | Moderate | Low (after scaling) | Quantitative | High |
| Human-based classification (e.g., serrated, lobed) | High | Moderate | None | No | High | None | Qualitative | Moderate (typically low) |
| Elliptic Fourier Transform | Low-Moderate (smoothing) | Low | High | No | Moderate | Low (typically PC axis 1) | Quantitative | High |
| Circularity | None | High | High | Yes | Low | High | Quantitative | Low |
| Shape index | None | High | High | Yes | Low | High | Quantitative | Low |
| Fractal dimension | None | High | High | Yes | Moderate | Low | Quantitative | High |
| Spectral/Approximate Entropy | None | High | High | No | High | None | Quantitative | High* |

\* Uniquely capable to detect localized features (anomalies) and quantify feature frequencies from signal curves

## METHODS

### Comparative framework

*ShapeComplexity*, designed for maximum performance and automation, requires no installation of dependencies but merely the local installation of an executable. The complete, open-source Rust-code (The Rust Team, 2025) is publicly available on GitHub (https://github.com/Thornbach/ShapeComplexity), ensuring transparency and reproducibility, while also encouraging community adaptation (Powers & Hampton 2019). Rust is a modern, high-performance language, capable of rapid processing large sets of high-resolution image data (∼1s per image). This Rust-based backend will be integrated into a user-friendly R-package (under development). The *ShapeComplexity* package we developed calculates aggregate shape metrics, leaf area, and our novel edge and macro-shape complexity metrics from single leaf images (Fig. 1). These (PNG) images require a transparent background and should ideally have the same resolution. Several tools are available online to prepare images easily, of which we used RMBG (https://huggingface.co/briaai/RMBG-2.0).

**Figure 1.**
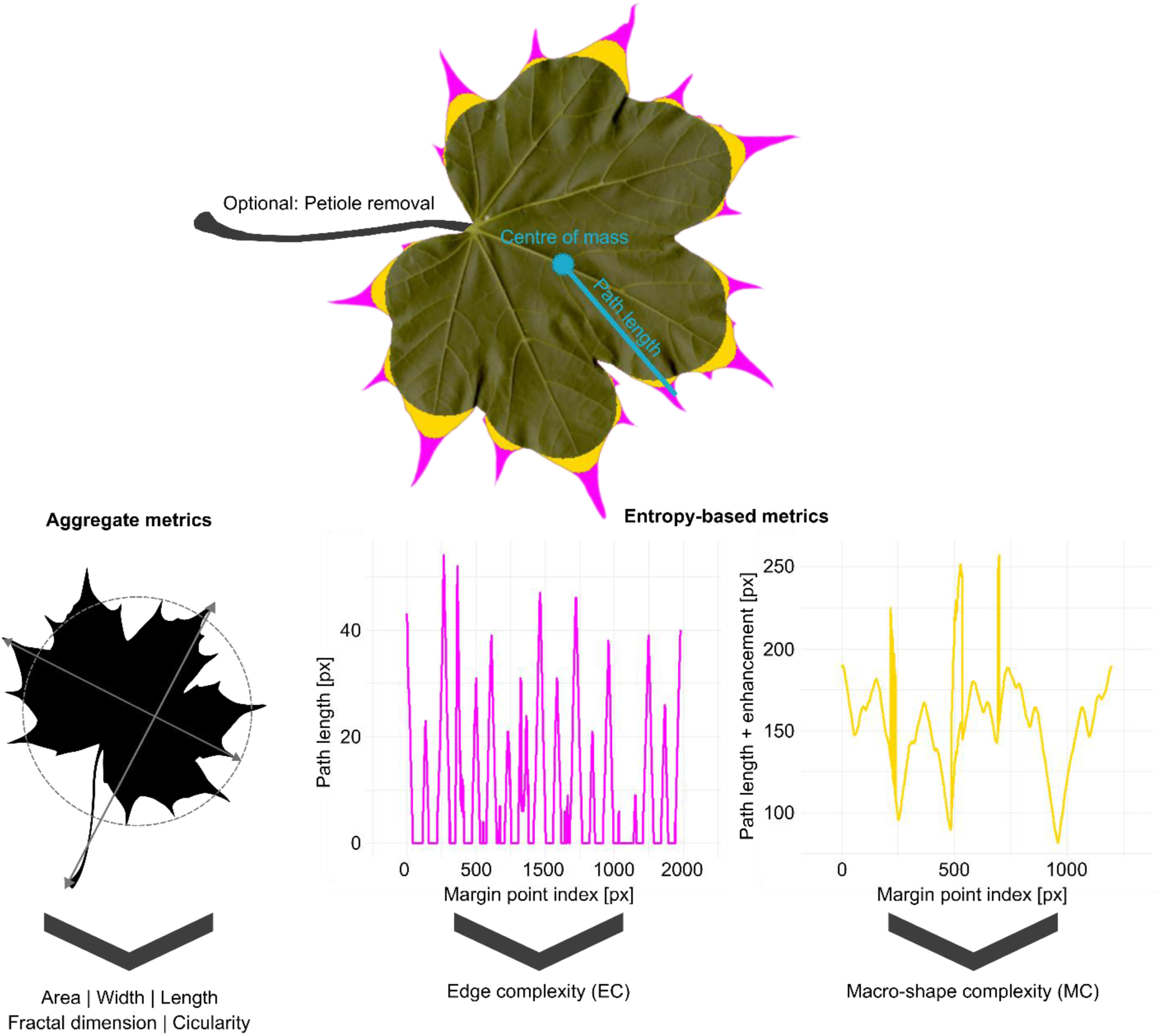
A simplified representation of the *ShapeComplexity* image analysis focused on the methods developed for calculating entropy-based leaf shape complexity metrics. Using images with a transparent background, an opening algorithm (i.e., inward erosion) is applied to retrieve two complementary layers for calculating EC (pink tips) and MC (yellow lobes). Subsequently, the intersections of paths ranging from the centre of mass to each consecutive contour pixel with the two layers are calculated, converting the layers into signal curves. Finally, the approximate entropy of the signal curve of the EC layer is calculated to quantify edge complexity and the spectral entropy of the signal curve of the MC layer is calculated to quantify macro-shape complexity. The output CSV file includes the proposed EC and MC metrics as well as fractal dimension and the widely used metrics circularity, shape index, and leaf area. The latter four of which do not distinguish (aggregate) edge and macro-shape complexity.

#### Theoretical foundation

Both EC and MC can be understood as empirical approximations of the Kolmogorov complexity, which is described as the length of the minimal description required to faithfully encode a given pattern (Kolmogorov 1968; Li & Vitányi 1997). A structurally simple shape, such as a perfect circle or an entire and smooth leaf margin, can be fully described by a short set of rules, yielding low Kolmogorov complexity. A toothed or deeply lobed outline are more difficult to describe, because each structural irregularity must be enumerated independently, increasing the required description length. Because Kolmogorov complexity is formally not computable, practical analyses rely on information-theoretic approximations (Cilibrasi & Vitanyi 2003). In *ShapeComplexity*, we implement two such approximations targeting independent spatial scales: approximate entropy for fine-scale edge features (EC) and spectral entropy for macro-scale shape variation (MC). Both are rooted in Shannon’s (Shannon 1948) foundational framework, which quantifies the expected information content of a probability distribution. Within *ShapeComplexity*, we apply it to the distribution of edge irregularities and shape frequencies respectively.

#### Aggregate metrics

For method validation and applications beyond the quantification of leaf complexity, *ShapeComplexity* also calculates circularity, leaf area, and the leaf shape index (the most widely used morphological traits) as well as fractal dimension. Leaf area is calculated based on the number of leaf (non-background) pixels and used to analyse the area-dependence of shape metrics and morpho space. For further analysis the pixel counts can be converted to metric units post-hoc through multiplication with the square of a scaling factor corresponding to the image resolution ((2.54/600)^2^ for scans with 600dpi resolution to obtain cm^2^). Circularity is calculated with the following formula: *4π * Area / Perimeter²*. A value of 1.0 indicates a perfect circle, while values approaching 0.0 indicate the opposite. The shape index (length-to-width ratio) is calculated as the longest Euclidean distance between outline pixels [*max(sqrt((x₂ - x₁)² + (y₂ - y₁)²)*] divided by the longest distance of its orthogonal. Fractal dimension is estimated using the box-counting method (Mandelbrot 1967): the contour is overlaid with grids of decreasing box size epsilon, the number of boxes intersecting the contour is counted at each scale, and the fractal dimension D is the negative slope of the log-log regression of box count on box size.

#### Preprocessing – morphological operations

The analytical strategy of *ShapeComplexity* rests on morphological image processing, which includes a set of operations that probe the geometric structure of a shape using a kernel (Serra 1982). The first operation is the erosion which shrinks a shape by removing pixels at its boundary. A pixel at position (x, y) is retained in the eroded image only if every pixel within the neighbourhood defined by the kernel is also non-transparent in the source image. Any pixel whose neighbourhood extends outside of the object is set to transparent. The result is an inward contraction (rather than smoothing) of the shape by the kernel radius, removing effectively all visible pixels smaller than the kernel. The second operation is the inverse, called dilation: a pixel is set to non-transparent if any pixel within its kernel neighbourhood is non-transparent in the provided source image. Applied to an eroded image, dilation expands the contracted shape back outward, restoring its interior while the features removed by erosion are not recovered. Erosion followed by dilation with the same kernel is called image opening. The net effect is the removal of all serrations, teeth, indentations and similar features whose narrowest dimension is smaller than the kernel diameter, while the overall shape of the object is largely preserved. Opening is the core decomposition tool of *ShapeComplexity*, with the pixels present in the source image but absent after opening constituting the edge features and the remaining pixels the macro-shape (Fig. 1). A circular kernel is used in any of the previous listed operations to ensure that the operation is isotropic and hence independent of orientation of the feature.

#### Preprocessing – adaptive kernel size

The kernel diameter controls which features are removed during opening: a small kernel removes only the finest teeth, while a large kernel removes progressively coarser features. To ensure that the first opening scales consistently across images at different resolutions and with different amounts of leaf fill in the canvas, we implement an adaptive kernel size based on the fraction of non-transparent pixels. The opening percentage *p(rho)* is interpolated linearly between a minimum

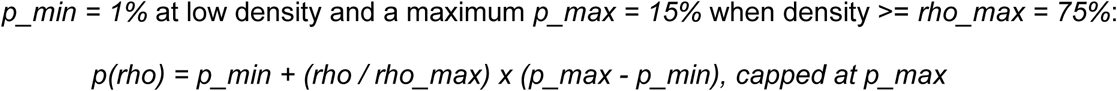

The kernel diameter is then:

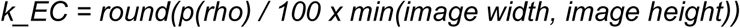

Using the shorter image dimension prevents excessively large kernels on non-square canvases. The result is a kernel that is always expressed as a fixed proportion of the leaf’s dimension on the picture itself rather than an absolute pixel count, making the method resolution independent.

#### Preprocessing – EC layer and MC layer separation

*ShapeComplexity* separates the shape into two spatially distinct layers by applying morphological opening twice in the sequence (Fig. 1). EC layer: The adaptive kernel *k_EC* is applied to the source image. Pixels that are non-transparent in the input but become transparent after opening constitute the EC layer. They represent the fine-scale edge features such as teeth and serrations whose narrowest dimension is smaller than *k_EC*. These pixels are internally marked in a working copy of the image to enable downstream geodesic calculations. MC layer: The MC layer is essentially derived by subtracting the EC layer from the original image, setting EC pixels transparent. A minor additional morphological erosion ensures that artifacts and fragments are removed, and the last remaining component is kept as the MC layer.

#### Preprocessing – lobe scaffold for structural element detection

To enable the identification of discrete structural elements such as lobes a morphological opening is applied to the MC layer using a dynamically computed kernel k_lobe that scales with the shape of the MC layer itself. The kernel size is derived from the shape index of the MC layer (*SI_MC*; the ratio of biological length to perpendicular width). Near circular shapes receive the largest kernel (default maximum: 30 % of the shorter MC dimension), while highly elongated shapes receive the smallest (default minimum: 5%), with linear interpolation:

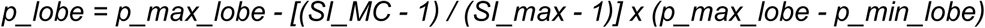

Where SI_max = 5.0 defines the upper elongation bound. The kernel diameter is:

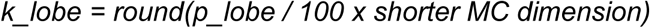

Pixels present in the MC layer image but removed by this aggressive opening are marked in a distinct colour. The resulting image, which will be called further the lobe scaffold, serves as a spatial reference for structural element assignment during geodesic analysis. The shape-adaptive kernel ensures that the lobe scaffold is meaningful for both circular species (e.g., *Populus tremula*, where the lobes project radially and are rather small requiring a proportionally larger kernel to isolate) and elongated species (e.g., Salix, where a smaller kernel suffices).

#### Outline linearisation and geodesic path calculation

The boundary of each image layer is extracted by scanning for edge pixels: non-transparent pixels that have at least one transparent pixel in their 8-connected neighbourhood. The resulting ordered sequence of contour pixels, traversed continuously around the perimeter, forms the signal scaffold onto which all path calculations are anchored. For each contour pixel, a path is computed from a reference point inside the leaf through the leaf interior. By default, the reference point is the centre of mass, which is the pixel-weighted centroid of all non-transparent pixels. The centre of mass is computed independently for the EC and MC images. An alternative emerge point (EP: the lowermost central pixel, representing the petiole insertion if the images are prepared accordingly) is available for studies where an anatomical anchor is preferred. The path computation proceeds in two stages:

1. Fast path (straight line): If the Euclidean straight line from reference point to contour pixel does not cross any transparent pixels and does not pass within two pixels of the leaf boundary at any intermediate point, it is used directly. The line is traced using Bresenham’s algorithm (Bresenham 1965). A two-pixel boundary zone is evaluated for all but the final quarter of the line length. This prevents silent inflation of edge-feature counts when the straight line grazes an unrelated tooth on the way to the target.

(2a) EC geodesic (Dijkstra with boundary penalty): When the straight line blocked or grazes the boundary zone, Dijkstra’s shortest-path algorithm (Dijkstra 1959) is applied on the marked image in which EC-layer pixels are opaque. A boundary penalty of 30 is added to the step cost of any pixel within two pixels of the leaf margin, making the boundary a near-prohibitive route. This steers the path through the leaf interior and prevents it from routing around a lobe by following the continuous EC strip along the margin, which would falsely inflate the edge-feature count for neighbouring teeth.

(2b) MC geodesic (breadth-first search): For the MC analysis, paths are computed on the MC layer image using breadth-first search, which treats all interior pixels as equally traversable. This approach yields the true unweighted geodesic distance, which represents the shortest hop count through the leaf tissue and hence the quantity for characterising macro-structural connectivity without edge-feature weighting.

#### EC signal: pink-pixel counting and filtering

For each EC contour pixel, the number of EC-layer pixels intersected by the geodesic path is counted. This yields a signal of length equal to the number of contour pixels, where each value represents the local prominence of the edge feature at that position, being zero for flat or entire margin regions and greater for teeth and serrations. Two noise-reduction filters are applied before entropy calculations: First, path intersections of <= 3 EC-layer pixels (default) are set to zero, removing digitisation artefacts. Second, an optional petiole filter can be applied as the petiole typically produces a large, isolated peak in the EC signal. Left uncorrected, this purely anatomical feature would inflate EC. The petiole is detected by a three step algorithm: (1) all contiguous runs of signal values exceeding a threshold of 1.0 are identified; (2) runs that straddle the start/end of the contour array (which indicates that contour tracing began mid-petiole) are merged into a single run using wrap-around detection; (3) only runs that contain at least one value at or above the 95^th^ percentile of the full signal are retained, and the longest qualifying run is designated the petiole sequence. If detected, the petiole pixels are excluded entirely from the EC signal, reducing the contour to the lamina only. This filter can be deactivated in the configuration if petioles have been removed through image preprocessing.

#### Structural element detection and harmonic enhancement

The raw geodesic path length signal is further modified by a harmonic enhancement procedure that amplifies the contribution of major lobes and other relevant and marked structures, increasing the sensitivity of spectral entropy to their number and size.

Structural element detection: for each contour pixel, the geodesic path is evaluated against the lobe scaffold image. A contiguous run of >= 15 contour pixels (default) whose geodesic paths each cross >= 5 scaffold pixels is identified as a valid structural chain. Each chain is characterised by its total length *L_seg* and its deepest point (the contour pixel whose point crosses the greatest number of scaffold pixels, indicating maximum penetration into the structural element).

Harmonic enhancement: The raw geodesic path length at each contour position within a valid chain is enhanced by superimposing a series of sine waves. The number of harmonics for a chain is proportional to its relative size:

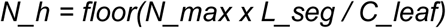

Where *N_max* = 12 (default) is the maximum number of harmonics and *C_leaf* is the total leaf circumference in contour pixels. Longer chains receive richer harmonic series, reflecting the greater spatial information content of larger structural elements. The enhancement intensity follows a linear position weight *W_pos* that equals 1.0 at the deepest point of the chain and falls to 0.0 at the chain boundaries. The final enhanced signal at contour position *i* within a chain is:

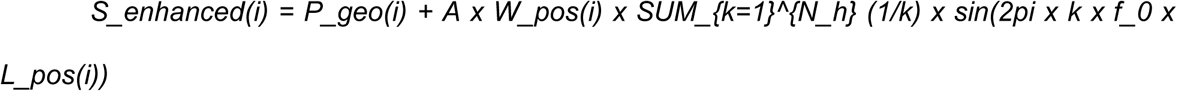

Where *P_geo(i)* is the raw geodesic path length; a = 2.0 is the harmonic strength multiplier (default), *f_0 = 1/C_leaf* is the base spatial frequency normalised by leaf circumference (ensuring scale invariance), and *L_pos(i)* is the cumulative contour position. The *1/k* amplitude weighting follows the harmonic series. Contour positions not belonging to any valid chain receive no enhancement.

#### Edge complexity (EC)

EC is calculated as the approximate entropy (ApEn; Pincus 1991) of the filtered EC-layer pixel count signal. Approximate entropy quantifies the likelihood that runs of similar values in a sequence remain similar when extended, with a regular, repeating signal having a low and an irregular, unpredictable signal having a high ApEn approximate entropy — analogous to high Kolmogorov complexity. Formally, *ApEn(m, r)* is defined as:

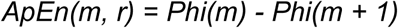

Where *Phi(m) = (1 / (N - m + 1)) x SUM_i ln(C_i^m / (N - m + 1))*, and *C_i^m* is the number of pattern windows of length *m* within tolerance *r* of the window starting at position *i* (using the Chebyshev / L-infinity norm). We use the template length *m = 2* (standard for short biological signals (Pincus 1991)) and a scale-adaptive tolerance *r = 0.2 x sigma*, where sigma is the standard deviation of the signal. This adaptive formulation ensures that the tolerance scales with signal amplitude, rendering EC independent of the absolute magnitude of edge features and enabling meaningful comparison across leaves of different sizes and resolution (Richman & Moorman 2000). A high EC indicates an irregular, unpredictable distribution of edge features characterizing, for instance, serrated or complex-toothed margin whose structure are mathematically more difficult to describe. A low EC indicated a regular or entire margin.

#### Macro-shape complexity (MC)

MC is calculated as the spectral entropy (Powell & Percival 1979) of the harmonically enhanced geodesic path length signal, processed through five steps:

1. Periodic Gaussian smoothing. The harmonically enhanced signal is smoothed with a periodic Gaussian filter (wrapping at contour boundaries) to reduce high-frequency digitisation noise. The window size is set to 1/8 of the signal length (minimum 3, maximum 21 points) and Gaussian sigma = 2.0 (default). Periodic boundary conditions treat the signal as a closed contour.
2. Power spectrum via FFT. A Fast Fourier Transform (Cooley & Tukey 1965) is applied to the smoothed signal after removal of the DC (mean) component. The power at each frequency *f_k* is computer as the squared magnitude of the Fourier coefficient: *P(f_k) = |X(f_k)|^2.* Only positive frequencies (k = 1 to N/2 – 1) are retained. The power spectrum is normalised so that *SUM P(f_k) = 1,* yielding a probability distribution over spatial frequencies.
3. Normalised Shannon entropy (Shannon 1948). The spectral Entropy is computed as *H_spectral = -SUM_k P(f_k) x log2(P(f_k))* and normalised by the maximum possible entropy *log2(N/2 - 1)* to yield values in [0, 1]. A simple leaf for example concentrates power in few frequency bins (low spectral entropy); a complex lobed leaf distributes power broadly (high spectral entropy).
4. Sigmoid noise suppression. For geometrically simple leaves, even minor digitisation noise may produce artifactually elevated raw spectral entropy, because noise energy is spread broadly across frequency bins. To suppress this effect, the raw entropy is scaled by a sigmoid function of the signal’s coefficient of variation *(CV = sigma_signal / mu_signal): S(CV) = 1 / (1 + exp(-k x (CV - c)))* where k = 20 (transition steepness; default) and c = 0.04 (transition centre; default). For signals with CV below c, S(CV) is near zero, suppressing noise-driven entropy in simple shapes. For *CV >= c, S(CV)* rises steeply toward 1.0, preserving the full entropy signal in genuinely complex shapes.
5. Structural complexity factor (Weber-Fechner scaling). The sigmoid-scaled entropy is further weighted by a logarithmic factor reflecting the number of valid structural chains *n* detected in the lobe scaffold *f(n) = ln(1 + n) / ln(1 + n_0)* where *n_0 = 10* is a reference value representing a highly complex leaf. This weighting follows the Weber-Fencher principle (Fechner 1860), which describes the psychophysical finding that perceived complexity increases sub-linearly with the number of structural elements. A leaf with no detected structural element (n = 0) receives the minimum baseline factor of 0.1; a leaf with n = 10 chains receives the maximum factor of 1.0. The final MC value is: *MC = H_spectral x S(CV) x f(n)*

### Statistical analysis

All statistical analyses were conducted in R (version 4.4.1) (R Core Development Team, 2024).

#### Case study 1: Linking leaf shape and leaf chemistry in Q. robur

To contextualize our framework, we compared the performance of our two novel complexity metrics (EC and MC) against four conventional morphological traits: circularity, leaf area, shape index, and fractal dimension. To explore the relationships between all six morphological traits, we first performed a principal component analysis (PCA) on the normalised (centred and scaled) data using the *prcomp* function from the base R-package *stats*. To then assess the association between these morphological traits and four chemical traits, we fitted a series of single-predictor mixed-effects models. For normally distributed responses we used linear mixed models, and for the proportionally scaled anthocyanin and nitrogen balance index data we used beta generalised linear mixed models (which use a logit-link function). All models included the tree identity as a random slope effect. Finally, we compared the predictive power of all models for each chemical trait using the Akaike Information Criterion (AIC) and their marginal (R²m) and conditional (R²c) explained variance to rank the predictive power of morphological traits.

#### Case study 2: Interspecific morphological separation

For our interspecific analysis, we quantified the same six morphological traits used in the first case study (EC, MC, circularity, leaf area, shape index, and fractal dimension) for a set of 902 leaf images from 13 tree species across nine genera (Söderkvist 2001). We conducted two distinct analyses. First, to explore the relationships between morphological traits, we performed a PCA on all six normalised (centered and scaled) traits using the *prcomp* function from the base R-package *stats*. Second, to evaluate the ability of different metric sets to create biologically intuitive clusters, we compared the morphospace spanned by our two novel complexity metrics (EC and MC) against the space spanned by leaf area and fractal dimension. The latter pairing was specifically chosen because our preceding Procrustes analysis identified it as the most complementary two-variable combination capturing the total morphological variation among aggregate metrics as per the interspecific PCA. For both morphospaces, the separation of genera was quantified based on pairwise overlap (Schoener’s *D* index) among genus-level Gaussian kernel density clusters (defined by the 95% confidence intervals), using functions of the R-package *ecospat* (Di Cola *et al*. 2017).

#### Case study 3: Human perception of complexity

To ground our quantitative metrics in a real-world benchmark, we tested how well they aligned with human visual perception. Using leaf silhouettes from the Swedish Leaf dataset (Söderkvist 2001), we conducted a survey with 90 participants via a custom R Shiny application. In the survey, participants were shown random pairs of leaves and asked to select the one they perceived as “more complex” (min: 10, max: 400, median: 51; user-choice). The final data comprised 7,759 individual ratings for 195 unique leaves.

We first assessed the consistency of the ratings by calculating Fleiss’ Kappa, a metric of agreement across raters for the same pairs, on a validation set of the 18 most frequently evaluated pairs (the top 10% of votes, cumulatively). This inter-rater reliability analysis on a validation set with mixed difficulty levels showed fair agreement (Fleiss’ Kappa = 0.38), confirming a consistent, non-random consensus among participants (see also Supporting Information – Method detail).

We then calculated a human perception score for each leaf, defined as the number of times it was chosen as more complex divided by the number of times it was presented to the survey participants (Sato 2009). To determine which metric best aligns with human perception, we assessed the pairwise relationship between each of the six morphological metrics and the human perception score using Pearson’s correlation coefficient. Finally, we performed a k-means clustering of combined EC and MC metrics and used Tukey’s HSD test to assess the association of the resulting clusters with the mean human perception score.

## RESULTS

### Case study 1: Intraspecific variation in leaf complexity and chemistry in *Q. robur*

The first two principal components of the six morphological metrics (EC, MC, fractal dimension, leaf area, circularity, and shape index) explained 75.1% of the overall shape variation. Our novel metrics, EC and MC, contributed strongly and independently to one component each (Fig. 2). The first principal component (41.9% variance) described a gradient in overall size and macro-scale complexity, with strong positive loadings from our MC metric, fractal dimension, and leaf area. The second component (33.2% variance) described fine-scale edge features and leaf elongation, with a strong positive loading for EC and negative loadings for circularity and leaf area. Despite directional alignment between the aggregate and novel metrics, both PCA axes loadings and quantitative correlation analysis (Supplementary Fig. S1, Fig. S2) showed a greater performance of MC and EC in quantifying leaf shapes. In this specific case of *Q. robur,* which has entire margins, the EC metric effectively captured the prominence and sharpness of the major lobes.

**Figure 2.**
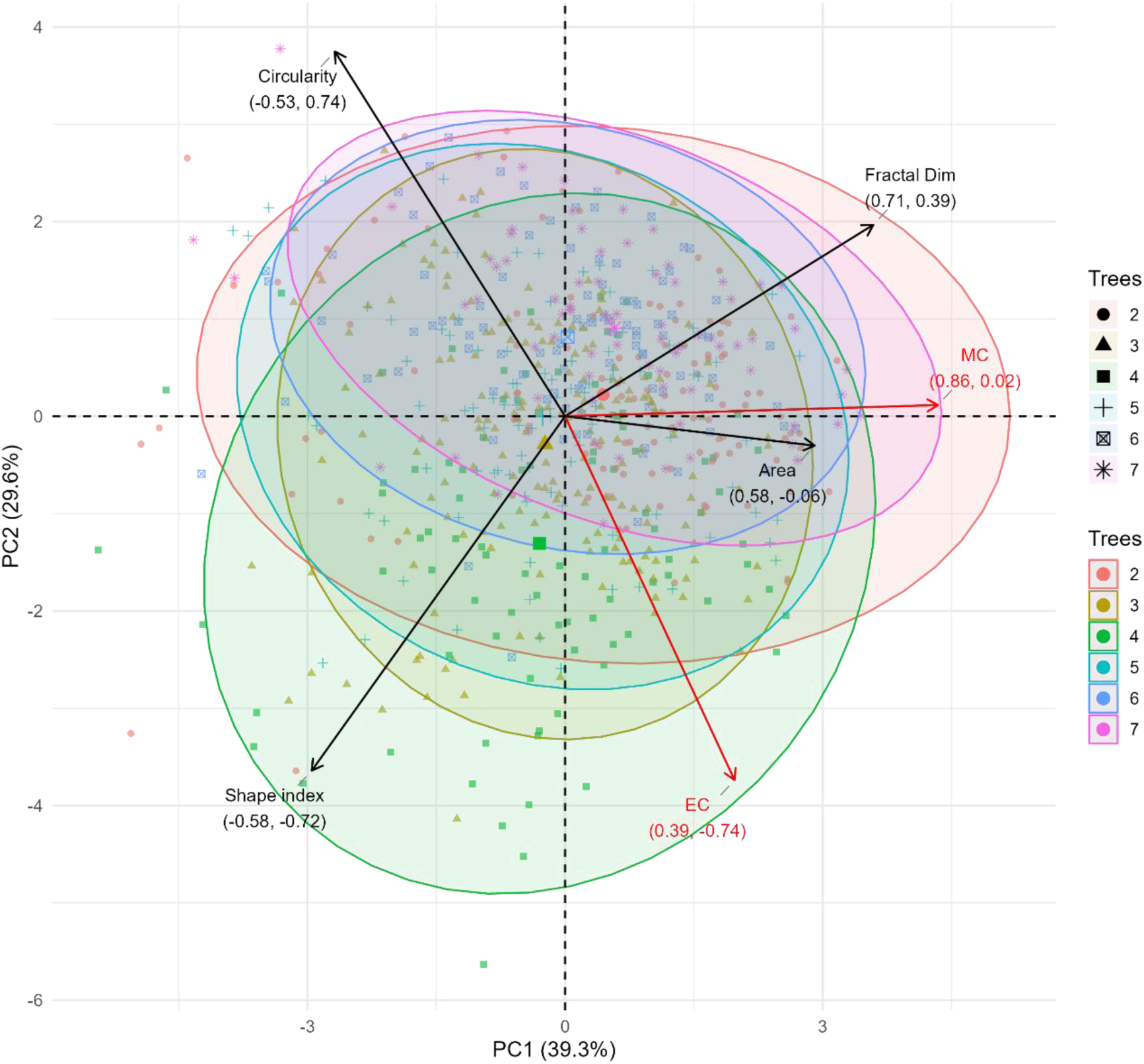
Intraspecific variation in leave shape complexity. Principal component analysis of the relationships between six morphological metrics and the distribution of 696 leaves of 6 *Quercus robur* trees. The analysis included two newly proposed (EC and MC, in red), one established but rarely applied (fractal dimension) and the most widely used (circularity, shape index and area) metrics of shape complexity. Arrows for the metrics represent variable loadings (% contribution). Ellipses indicate the 95% confidence intervals for each tree.

Associations of leaf morphology and chemistry revealed two clear patterns. First, the random factor of sun/shade nested in season and tree identity (of which the former two factors were least relevant) was the primary driver of variation in all models, with the full models explaining between 22.2% and 71.8% of the variance in chemical traits (R²c; Supplementary Table S2). After accounting for this strong individual-level variation, EC emerged as the best predictor for flavonoids and anthocyanins, while area was the top predictor for chlorophyll and nitrogen balance index among shape complexity metrics according to model AICs (Fig. 3). EC explained 4.7% of the variance for flavonoids and 4.0% for anthocyanins (R²m). Circularity was a weaker predictor across all traits, explaining between 1.4% and 2.4% of the variance.

**Figure 3.**
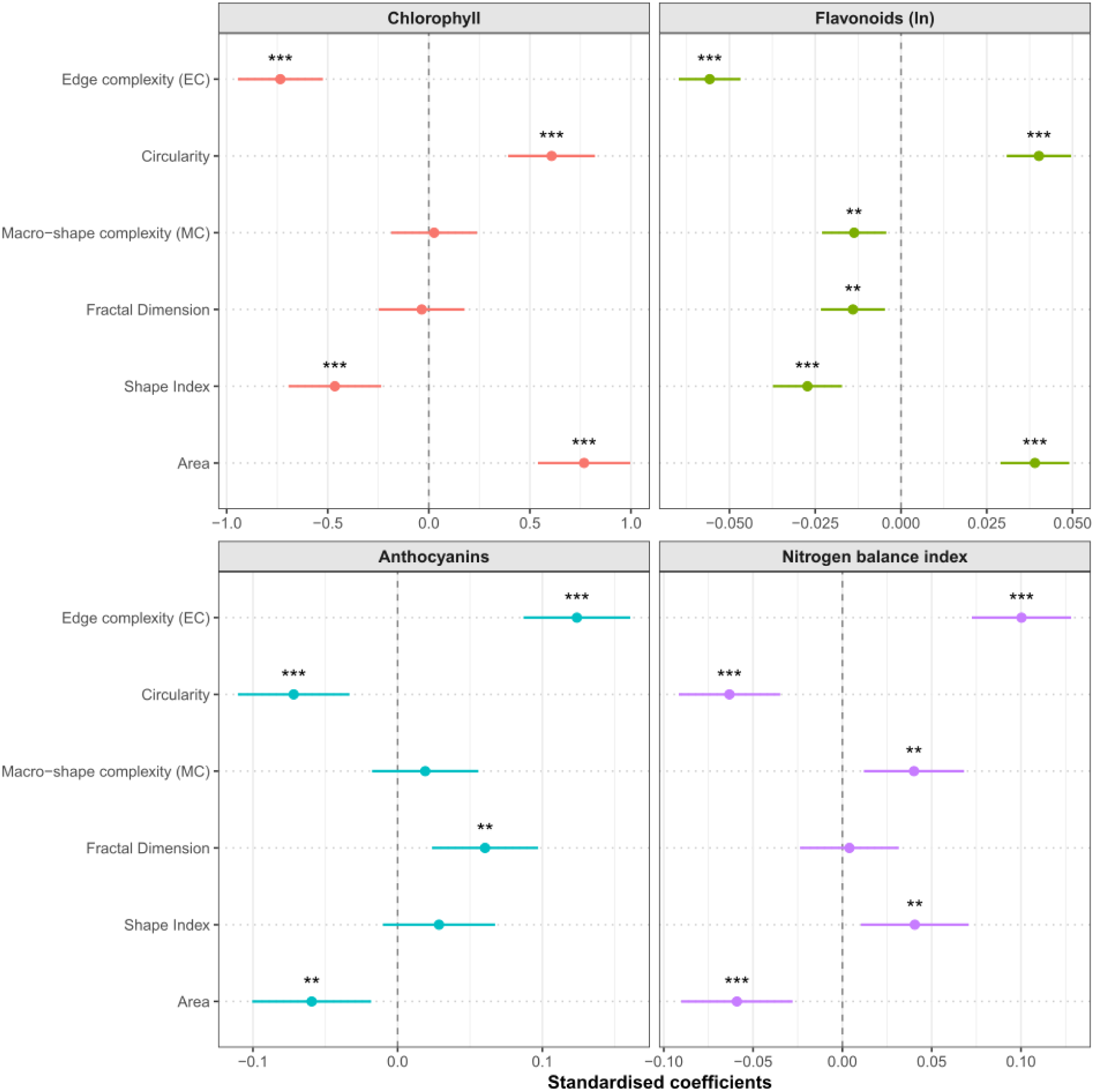
Relationships of complexity metrics with intraspecific variation in chemical leave traits. Results from single mixed-effects models of the relationships of six shape complexity metrics with chlorophyll content, flavonoid (log_10_-transformed) content, anthocyanin content, and the nitrogen balance index across 696 leaves of 6 *Quercus robur* trees. Leaf area is an important morphological trait but not a proxy of shape complexity, here considered only for reasons of comparability with the morphospace analysis. All models included a random slope effect for light regime nested in tree identity. Points and bars are standardised coefficients and their 95% confidence intervals, respectively. Asterisks indicate significance at p < 0.05*, p < 0.01**, p < 0.001***.

### Case study 2: Species classification using interspecific leaf shape variation

The first two principal components of the same six morphological metrics explained 74.4% of the overall interspecific shape variation. In contrast to the intraspecific analysis, this PCA demonstrated that our novel metrics, EC and MC, define the primary component of shape complexity in concert, with area most effectively complementing as per the secondary component. The first principal component (39.6% variance), which separated simple from complex leaves, was similarly strongly driven by fractal dimension, EC, and MC (Fig. 4). The second principal component (34.8% variance) represented a distinct axis of leaf size and elongation and was primarily driven by the aggregate metrics area, circularity, and shape index.

**Figure 4.**
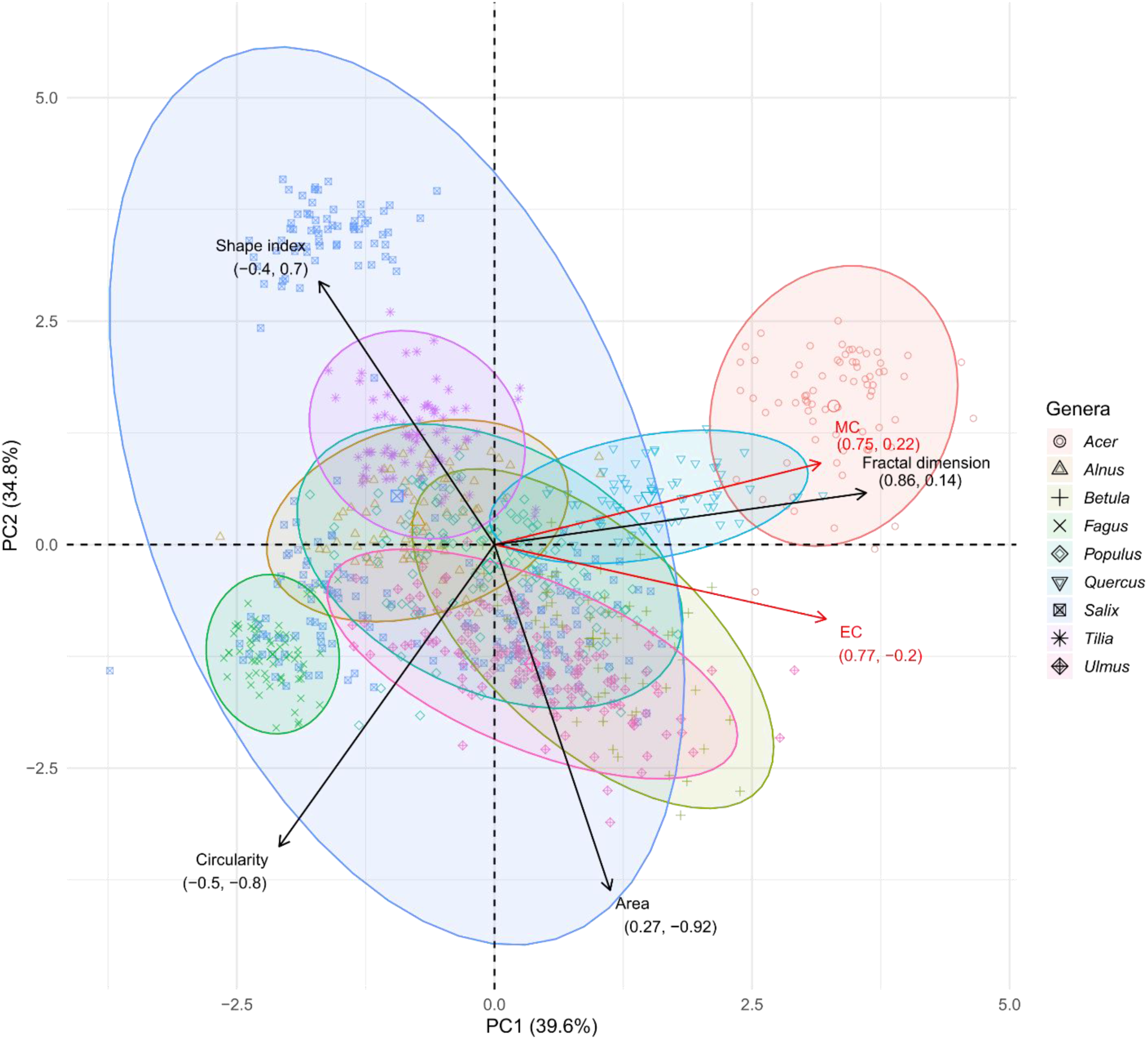
Interspecific variation in leave shape complexity. Results of a principal component analysis of the relationships between six morphological metrics and the distribution of 902 leaves from 9 tree genera sampled across Sweden. The analysis included two newly proposed (EC and MC, in red), one established but rarely applied (fractal dimension) and the most widely used (circularity, shape index and area) metrics of shape complexity. Loadings (% contribution) of the six metrics are indicated by vectors and the numbers below their labels. Ellipses indicate the 95% confidence intervals for each genus. Note the high redundancy (orthogonal vectors) of aggregate metrics as well as the overall similarity of vectors here and for intraspecific variation across leaves of *Quercus robur* individuals (Fig. 2).

When comparing the ability of different morphospaces to separate the nine genera, both the entropy-based space (EC and MC) and the shape variation characterised by leaf area and fractal dimension yielded highly distinct species-level clusters. The entropy-based space positioned species in a way that was more consistent with their visible leaf shape complexity compared to the aggregate metrics (Fig. 5A, B) and had a slightly lower species overlap (Schoener’s *D* = 0.11 < 0.13, Fig. 5C, D). For instance, whereas the latter confused the deeply lobed leaves of *Quercus* with the simple, round leaves of *Populus* species (*D* = 0.32), our entropy-based metrics correctly separated these morphologically distinct species (*D* = 0.00). Conversely, our metrics showed high overlap between the visually similar *Salix* and *Ulmus* species (*D* = 0.72), which the area and fractal dimension combination failed to capture at this level (*D* = 0.40).

**Figure 5.**
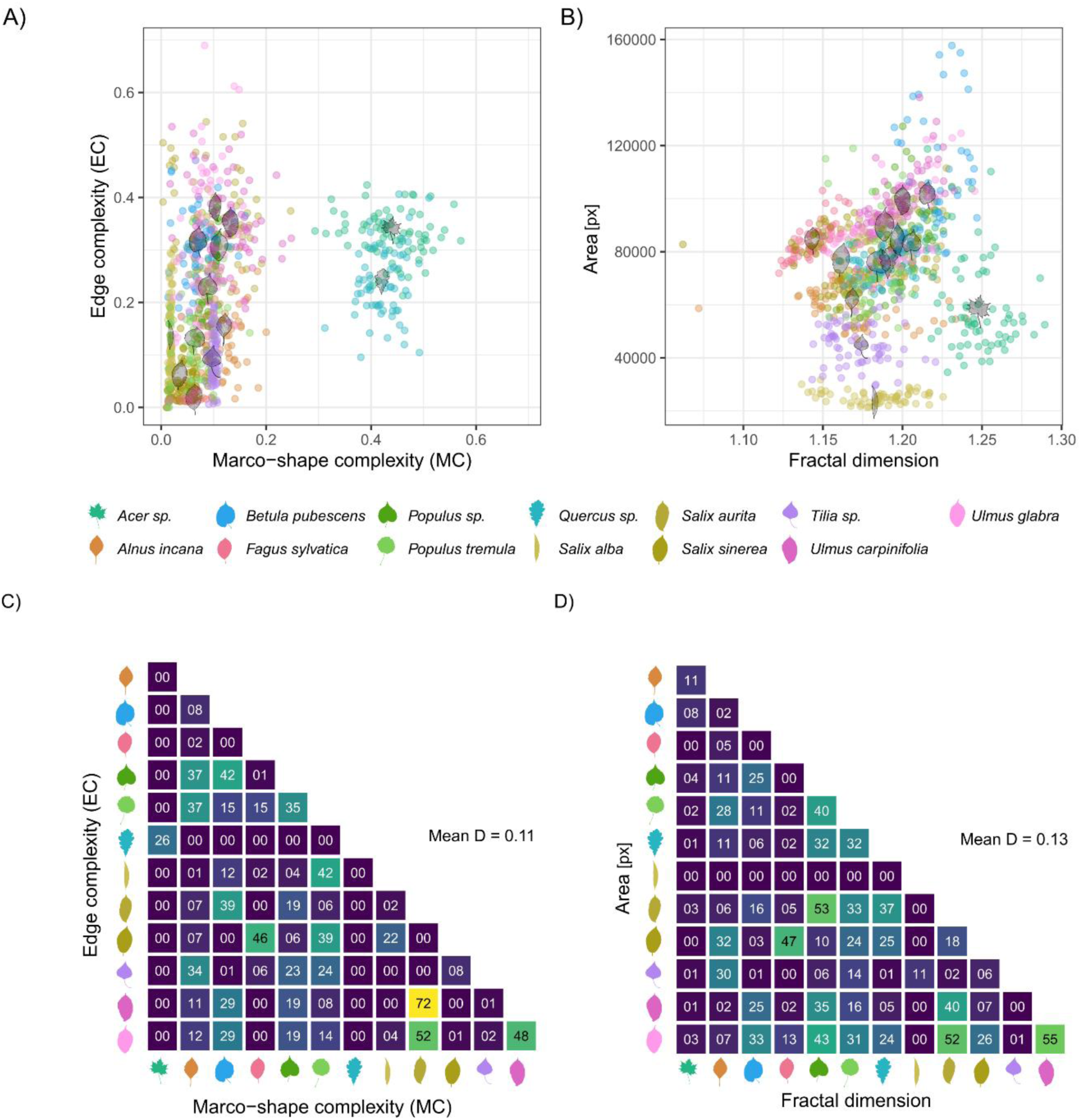
Species’ leave shape similarity as per entropy-based on established complexity metrics. Comparison of the leaf complexity of 902 leaves of 13 species for 9 tree genera across Sweden. (A) Leaf complexity variation defined by approximate entropy (EC, edge complexity) and spectral entropy (MC, macro-shape complexity) in comparison to (B) the morphospace spanned by area and fractal dimension. Each point represents an individual leaf, coloured by species. Black silhouettes show a representative leaf shape for each species, positioned at the genus centroid. Matrices in the bottom panel show the pairwise species overlap (Schoener’s *D* × 100) for our novel entropy-based metrics (C) and for the two metrics leaf area and fractal dimension (D).

### Case study 3: Human perception of complexity

Our survey of human perception revealed that the calculated scores were strongly aligned with our entropy-based metrics. The EC, which quantifies fine-scale edge complexity, was by far the single best predictor (*r* = 0.76, p < 0.001). Fractal dimension was the second strongest (*r* = 0.62, p < 0.001) and our MC metric for overall shape complexity was the third-strongest predictor (*r* = 0.52, p < 0.001) for complexity as perceived by humans (Fig. 6A).

**Figure 6.**
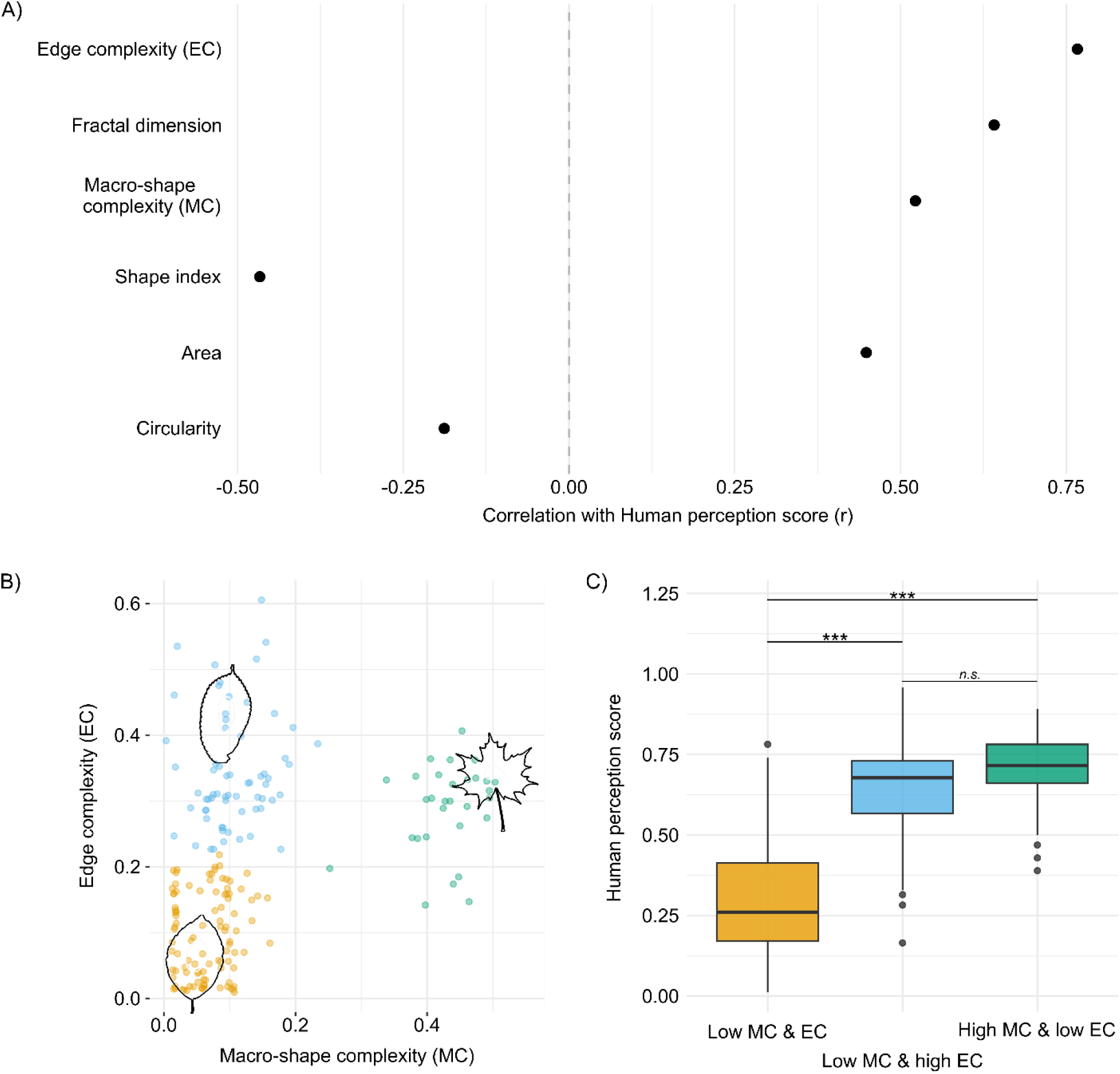
Alignment of leaf complexity metrics and leaf complexity as perceived by humans. (A) Correlation (Pearson’s *r*) of six morphological metrics with the Human Perception score. (B) Complexity variation as defined by leaf edge and leaf macro-shape complexity, showing three k-means clusters. Representative leaf silhouettes are plotted at the centre of each cluster. (C) Boxplots show the Human Perception scores for the three complexity clusters (“High edge complexity” but “Low macro-shape complexity”, “Low edge complexity” but “High macro-shape complexity”, “Low edge complexity” and “Low macro-shape complexity”). Tukey’s HSD test results are indicated as: *** p < 0.001, n.s. = not significant.

An ANOVA comparing the three k-means clusters based on EC and MC corresponded to perceptually distinct categories of complexity (F(2, 192) = 115.8, p < 0.001). The “Low EC and low MC” cluster (n = 92) was rated as significantly less complex than two other groups (p < 0.001 for both) (Fig. 6B, C). However, there was no significant difference in perceived complexity between the “High EC and low MC” group (n = 75), characterised by complex edge structures but simple macro-shapes, and the “Low EC and high MC” group (n = 28), characterised by minor edge structures but complex macro-shapes (p = 0.36). This result demonstrates that human perception recognizes two similarly powerful pathways to high complexity: one driven by intricate edge detail (captured by EC) and another by complex overall shape (captured by MC).

## DISCUSSION

Together, our three use cases demonstrate the major predictive power, unique explanatory contribution, functional plausibility, and versatile application of deconstructing leaf shape into edge complexity (EC) and macro-shape complexity (MC). Across both our intra- and interspecific analyses, these new metrics consistently captured the primary axes of shape variation. Within *Q. robur*, entropy-based metrics also generally had the highest predictive power for chemical traits. Interspecifically, EC and MC outperformed the most relevant aggregate metrics in classifying species. In fact, while the latter notably failed to account for apparent edge complexity, entropy-based metrics provided not only an empirically better species-classification but also more intuitive clustering of similar shapes among species. Towards understanding visual cues used for classifying complexity, our case study on human perception revealed that both EC and MC had far greater effects on perceived complexity than aggregate metrics. Implemented in our package *ShapeComplexity*, these metrics now enable automated and nuanced assessments of shape variation across spatial and temporal scales, well beyond leaves—as demonstrated by our algae use-case—with broad applications in ecology and evolution.

### Leaf edge complexity as a proxy for leaf chemistry

Our results on intraspecific variation in leaf shape and chemical traits in *Q. robur* highlight the greater functional relevance of edge complexity compared with macro-shape complexity and aggregate metrics for this heterophyllous species at the local scale. Notably, as we minimized the impact of environmental conditions by selecting trees in a homogenous forest patch, the overwhelming importance of tree identity suggests that genetic differences rather than shared responses to microclimatic conditions and development underpin decoupled variation in leaf macro-shape complexity and chemical traits at this scale. Thus, while greater coupling can be expected across broader environmental gradients, at the local scale MC was only weakly positively related to nitrogen balance index and negatively related with flavonoids.

The nuanced differences captured by EC explained more than any other complexity metrics and a relatively large share of the chemical trait variation independent of tree identity. Specifically, EC was positively associated with anthocyanins and the nitrogen balance index, and negatively related to chlorophyll and flavonoid content. These relationships are consistent with evidence that increased edge complexity enhances gas exchange, transpiration, and nutrient transport (Royer & Wilf 2006; Wilf 1997), and that more dissected leaves exhibit higher anthocyanin and lower chlorophyll content, reflecting increased light exposure and localized photoprotective responses (Agati & Tattini 2010; Givnish 1979; Niinemets 2007). In addition, our results support previous evidence that leaves with pronounced lobes or serrations experience intensified light and thermal gradients at their margins, promoting localized anthocyanin synthesis and elevated nitrogen balance indices (Sack & Holbrook 2006; Vogel 2009). The ability of our entropy-based metrics to explain chemical variation beyond that captured by aggregate traits thus promises great potential for using morphological proxies to characterize the plant chemical spectrum also within species and individuals (Pérez-Harguindeguy *et al*. 2013; Wright *et al*. 2004). This provides a reproducible, non-invasive tool for assessing internal physiological and stress-related responses from simple leaf images (Agati *et al*. 2012; Fiorani & Schurr 2013).

### Addressing taxonomical and functional shape variation across species

Our entropy-based framework provided a more biologically intuitive and functionally meaningful representation of interspecific leaf shape variation compared to the most important and complementary aggregate metrics (i.e., area and fractal dimension). The morphospace spanned by the latter produced a poorer overall separation and yielded counter-intuitive groupings, such as conflating the deeply lobed leaves of *Quercus* with the simple, round leaves of *Populus* species. In contrast, the entropy-based morphospace produced distinct clusters that aligned with visible morphology. For example, it successfully captured the visual similarity of *Salix aurita* and *Ulmus carpinifolia* with a high degree of overlap, a grouping that the morphospace spanned by area and fractal dimension represented only moderately. This demonstrates that our framework, by integrating edge and macro-shape complexity, more effectively identifies leaves that are functionally similar and present analogous visual cues to herbivores (Rausher 1978).

### Perceptual validation of the framework

Our survey of perceived complexity strongly underscored the more intuitive entropy-based clustering and its association with visual cues. EC, quantifying fine-scale marginal irregularity, was the single strongest predictor of human-perceived complexity. Furthermore, the combination of EC and MC defined perceptually distinct complexity classes, confirming that humans weigh edge and macro-shape similarly when evaluating leaf form, despite the very minor impact of the inherently small edge features on aggregate metrics. The strong agreement between our entropy-based metrics and human perception suggests that our framework successfully captures dimensions of leaf form that are ecologically relevant. By providing a quantitative proxy for what “looks similar”, our metrics offer a reproducible, scalable and mechanistically tailored method for exploring the functional significance of morphological diversity, including but not limited to the ecology of pollinator choice and herbivore host-plant selection (Rausher 1978; Schoonhoven *et al*. 2010).

### Scope and limitations

A key strength of the proposed entropy-based complexity metrics is their inherent independence from leaf area, which renders most aggregate metrics highly redundant and strongly limits their information content (Fig. 4). Particularly across species the widely used metrics shape index and circularity strongly covary with, but are also outperformed by, leaf area in characterizing the overall shape variation of leaves (Figs. S1, S2).

Our method is designed for broad applicability, and its path-based approach is well-suited to handle complex forms including compound or fenestrated leaves (e.g., *Monstera*). Beyond botany, the mathematical principles of EC and MC are applicable to any closed biological contour (S3). Finally, the complete methods and workflow are implemented in our open-source package, *ShapeComplexity*, ensuring transparency and reproducibility, while encouraging community adoption. This is a critical aspect of advancing ecological methodology (Powers & Hampton 2019).

To maximize information content and reduce measurement errors *ShapeComplexity* benefits from high-quality input images of uncurled, non-overlapping objects with backgrounds removed. Users should select objects without significant damage or decrease the sensitivity in calculations of EC (i.e., decrease harmonic enhancement; see also Table S1) for analysis of general shape complexity. However, a high sensitivity can also be advantageous for analysis that aim to localize and assess herbivore damage or ontogenetic impacts (e.g., by galls or environmental stress), or simply for quality control and preselection of intact input images.

### Broader applications

The applications of our proposed entropy-metrics (MC and EC) extend well beyond the provided case studies. In plant science, these novel metrics and their automated assessment are an important step towards non-invasive spatially and temporally explicit monitoring of environmental stress (Gill *et al*. 2022). Similarly, researchers can readily use *ShapeComplexity* and the implemented metrics on fossilised leaves to quantitatively assess the evolution of different shape dimensions and their links to past climatic events (Royer *et al*. 2005). Our framework for decoupling EC and MC and the implementation of entropy-based metrics thus promise to unlock the potential of growing digital image archives of leaves, including images from herbaria and fossils, but are also readily applicable to shapes of algae, bacteria, pollen, and beyond (see Fig. 7).

**Figure 7.**
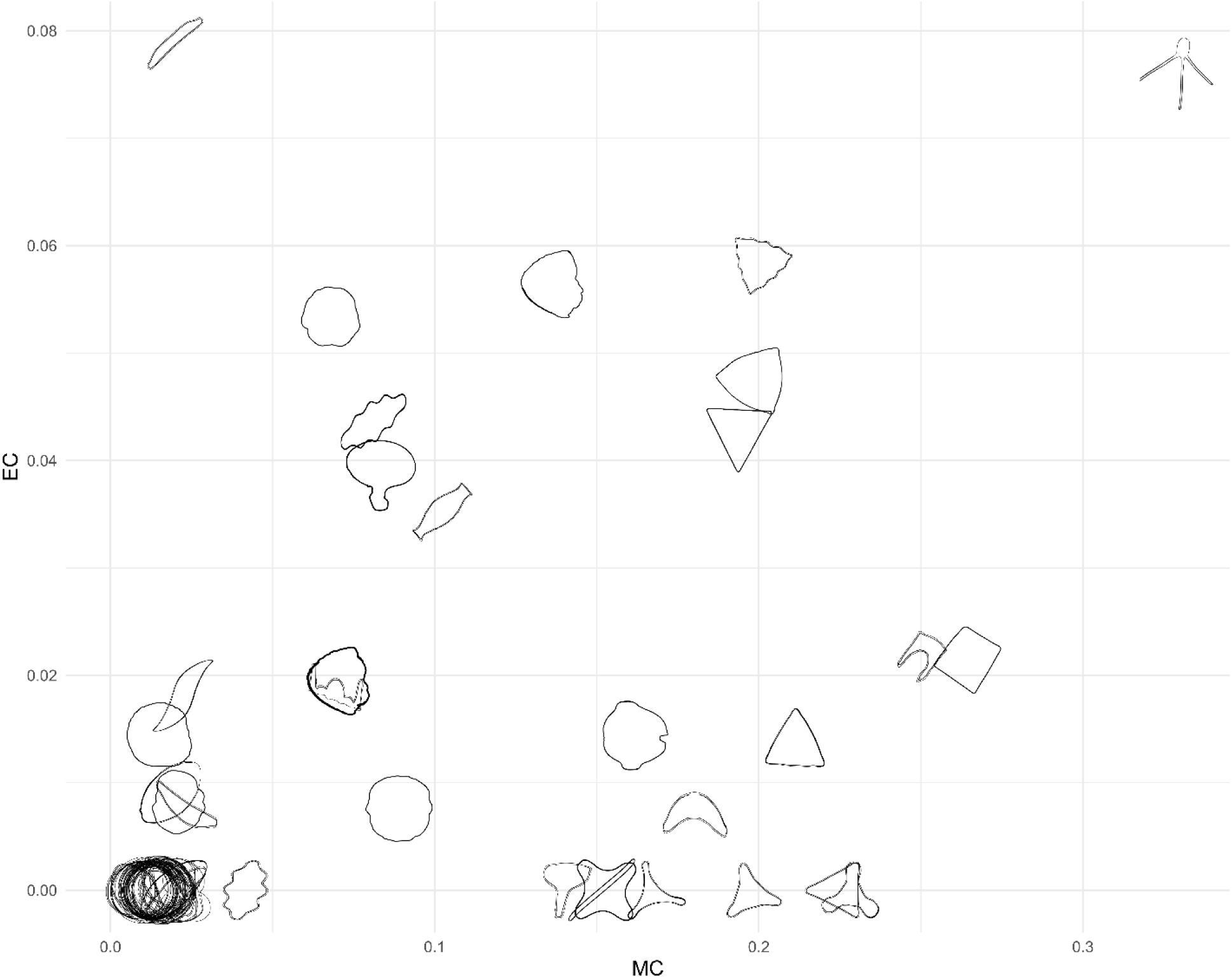
Algae use-case illustrating the transferability of the edge and macro-shape complexity framework. Application of the entropy-based framework to diatom images, quantifying edge and macro-shape complexity (EC and MC) metrics on diatom images (from artwork – microscopic images - by Alex Hyde). To improve visual clarity amid the abundance of circular shapes clustering in the bottom-left corner (i.e., very low EC and MC), only a random subset 50% of shapes (n = 134) was displayed.

### Conclusions

Established metrics have struggled to capture the multifaceted nature of leaf shape complexity, often conflating size, edge structure, and overall form into a single, ambiguous metric. Manual classifications and landmark-based approaches have remained the only means towards a unifying and explicit incorporation of these main dimensions, yet they are subjective and inherently difficult to reproduce. Here, we present a framework that decomposes leaf shapes into two highly complementary and functionally relevant dimensions: edge complexity (EC) and macro-shape complexity (MC). This two-component approach moves beyond simple technical refinement. It provides strong, mechanistically tailored predictors for leaf chemistry and fine-scale edge complexity, while also offering nuanced descriptors of functional adaptation and interspecific macro-shape variation, a distinction that, as we show, aligns closely with human perception. Implemented in *ShapeComplexity*, our method provides a rapid and scalable means to quantify structural complexity across taxonomic, spatial, and temporal scales, enabling rigorous integration of leaf shape variation into trait-based, ecological, and evolutionary analyses.

## Author Contributions

T.T., N.F. and S.P. conceived the study. T.T. performed all analysis. T.T. and S.P. wrote the first draft of the manuscript with support from N.F., A.R., M.M.A., R.D., K.T. and L.S. contributed data. T.T. and K. T. conceived and realised the survey. All authors contributed substantially to revisions.

## Acknowledgements

We are grateful to Kathina Muessig, Vicky Tough, Mona Schreiber, Lara Lohmann and Alex Stylianou for their help with collecting *Quercus robur* leaves as well as to the faculty members and students who participated in the survey. T.T., A.R., R.J., L.S., N.F. and S.P. acknowledge support by the LOEWE research initiative of the State of Hesse, Germany, through the Ministry of Science and Arts (HMWK), as part of the LOEWE research cluster Tree-M (LOEWE/2/15/519/03/08.001(0002)/88).

## Ethics statement

Ethical approval was not required by the ethics committee of our institution as the study was classified as a “project with no or marginal risk”. Our human perception assessment was conducted in full compliance with the principles of the Declaration of Helsinki. All participants were over 18 years old and were informed about the purpose of the study prior to participation. Participation was voluntary and could be withdrawn at any time. To ensure anonymity, every participant was assigned a unique survey code instead of names. Participants were informed that by returning the completed survey, they provided consent for their anonymised data to be used in analyses.

## Open Research statement

The Rust code for calculating the leaf morphometrics, used in this article, is publicly available on GitHub (https://github.com/Thornbach/ShapeComplexity) and permanently archived at Zenodo (DOI: 10.5281/zenodo.22113804; version 1.0.0 was used for the analyses presented here). All supplementary data and analysis code, including the human perception survey data (Supporting Information 2-5), are archived at Dryad (DOI: 10.5061/dryad.83bk3jb6m).

