## Supplemental File 1 for "Introducing entropy-based metrics for quantifying edge- and macro-shape complexity in leaves and beyond"

**Content:**

Method detail

Table S1-2

Figs. S1-2

**Method detail**

***Sensitivity analysis for entropy-based metrics***

Our sensitivity analysis focused on five core algorithmic parameters that govern the key stages of morphological decomposition and complexity quantification. For each parameter, we tested a range of values (low, default, and high) to assess the stability of the final complexity rankings across the entire Swedish Leaf dataset (excluding compound leaves) as quantified by Spearman's rank correlation (ρ).

Overall sensitivity: All Spearman's ρ values exceed 0.92, demonstrating that the rank-order of leaf complexity is remarkably stable. The biological interpretation of which leaves are complex is not sensitive to minor or even moderate adjustments of the core parameters. Modular independence of complexity metrics: The perfect correlation (ρ = 1.00) for EC in parameters 2-5 (Table S1) is particularly strong. Differences in the strength of correlation coefficients for EC and MC further highlight that the logical and mathematical independence of parameters at different stages of the entropy analysis.

The R-code and data for this analysis can be found in the file *SuppInfo_2.zip*.

**TABLE S1** Sensitivity analysis of parameter ranges. Correlation (Spearman’s ρ) of estimates of macro-shape complexity (MC) and edge complexity (EC) calculated with default parameters versus minimum and maximum parameter settings.

| Parameter | Parameter setting | MC (ρ) | EC (ρ) |
| --- | --- | --- | --- |
| 1. Adaptive Max Opening | Low (10%) vs. Default (15%) | 0.94 | 0.92 |
|  | High (20%) vs. Default (15%) | 0.92 | 0.93 |
| 2. TF Max Opening (Lobes) | Low (20%) vs. Default (30%) | 0.97 | 1.00 |
|  | High (40%) vs. Default (30%) | 0.97 | 1.00 |
| 3. TF Pixel Threshold | Low (3px) vs. Default (5px) | 0.99 | 1.00 |
|  | High (7px) vs. Default (5px) | 1.00 | 1.00 |
| 4. Harmonic Chain Length | Low (10) vs. Default (15) | 1.00 | 1.00 |
|  | High (20) vs. Default (15) | 1.00 | 1.00 |
| 5. Sigmoid c Entropy | Low (0.02) vs. Default (0.03) | 1.00 | 1.00 |
|  | High (0.04) vs. Default (0.03) | 1.00 | 1.00 |

***Shape complexity rating analysis***

To validate the human perception data, we performed an inter-rater reliability analysis on a subset of "validation" pairs that were rated by multiple participants. The goal was to quantify the level of agreement between raters, correcting for the probability that they might agree by chance. Fleiss' Kappa is the standard metric for this type of analysis, involving multiple raters and categorical choices. However, our study design presented a key challenge: each item (leaf pair) was rated by a different number of participants. Standard implementations of Fleiss' Kappa in popular R-packages (e.g., *irr*, *DescTools*) are designed for datasets with a fixed number of raters per item and were found to produce erroneous results with our data structure. To ensure a correct and robust calculation, we therefore implemented Fleiss' Kappa manually, following the original formulas. This approach handles a variable number of raters per item and provides an accurate measure of inter-rater agreement for our specific dataset.

The analysis pipeline consists of three main stages:

1. Data preparation: Loading raw vote data and leaf metrics, standardizing leaf IDs across datasets, and cleaning the data.
2. Creation of a standardised rating matrix: For each validation pair, we robustly identified the two leaves presented and transformed the raw rating into a standardised "Choice 1" vs. "Choice 2" format. This data was then aggregated into a summary matrix where each row represents a unique leaf pair and columns represent the counts for each choice.

3	Manual calculation of Fleiss' Kappa: Using the summary matrix, we calculated the final Kappa statistic by applying its component formulas directly: *κ = (P̄ - Pₑ) / (1 - Pₑ)*. Here *Pᵢ* represents the proportion of agreeing pairs of raters for each item, *P̄* the mean of all *Pᵢ* values (overall observed agreement) and *Pₑ* the proportion of agreement expected by chance. The R-code and data for this analysis can be found in the file *SuppInfo_5.zip*.

**TABLE S2** Generalised single linear mixed effects models of the relationships between morphological and chemical traits. In addition, p-values from *DHARMa* (Hartig *et al.* 2024) tests for residual distribution, over- or underdispersion, and excess outliers, respectively, are provided, where low values (p < 0.05, in bold) indicate potential model misspecification. Bold values of marginal explained variance (R^2^m) highlight the strongest predictor per chemical trait. Flavanol content has been log-transformed. The corresponding code can be found in *SuppInfo_S3.zip*.

| GLM stats | | | | | | | | | DHARMa stats | | |
| --- | --- | --- | --- | --- | --- | --- | --- | --- | --- | --- | --- |
| Response | Predictor | Coef. | SEM | z/t | P | AIC | R²m | R²c | Dist. | Disp. | Outlier |
| Anthocyanins | EC | 1.24e^-01^ | 1.88e^-02^ | 6.59 | 4.44e^-11^ | -2872 | **0.04** | 0.26 | **3.69e^-22^** | 0.26 | 0.556 |
| Anthocyanins | Area | -7.57e^-02^ | 1.85e^-02^ | -4.09 | 4.38e^-05^ | -2846 | 0.02 | 0.25 | **2.47e^-23^** | 0.22 | 0.188 |
| Anthocyanins | Circularity | -7.18e^-02^ | 1.97e^-02^ | -3.65 | 2.67e^-04^ | -2843 | 0.01 | 0.23 | **1.24e^-23^** | 0.14 | 0.307 |
| Anthocyanins | Fractal dimension | 6.04e^-02^ | 1.87e^-02^ | 3.22 | 1.26e^-03^ | -2840 | 0.01 | 0.23 | **1.21e^-25^** | 0.26 | 0.556 |
| Anthocyanins | Shape index | 2.86e^-02^ | 1.98e^-02^ | 1.44 | 1.49e^-01^ | -2831 | 0.00 | 0.23 | **2.17e^-27^** | 0.18 | 0.769 |
| Anthocyanins | MC | 1.91e^-02^ | 1.87e^-02^ | 1.02 | 3.08e^-01^ | -2830 | 0.00 | 0.22 | **4.17e^-25^** | 0.16 | 0.769 |
| Chlorophyll | Area | 1.02e^+00^ | 1.07e^-01^ | 9.55 | 1.28e^-21^ | 8246 | **0.04** | 0.49 | **1.62e^-01^** | 0.89 | 0.005 |
| Chlorophyll | EC | -7.35e^-01^ | 1.07e^-01^ | -6.85 | 7.55e^-12^ | 8278 | 0.02 | 0.44 | **8.74e^-02^** | 0.99 | **0.018** |
| Chlorophyll | Circularity | 6.07e^-01^ | 1.10e^-01^ | 5.54 | 3.02e^-08^ | 8293 | 0.01 | 0.42 | **2.51e^-01^** | 1.00 | **0.005** |
| Chlorophyll | Shape index | -4.65e^-01^ | 1.17e^-01^ | -3.98 | 7.00e^-05^ | 8310 | 0.01 | 0.43 | **1.40e^-01^** | 0.98 | **0.005** |
| Chlorophyll | Fractal dimension | -3.54e^-02^ | 1.08e^-01^ | -0.33 | 7.44e^-01^ | 8321 | 0.00 | 0.42 | **1.12e^-01^** | 0.97 | **0.005** |
| Chlorophyll | MC | 2.62e^-02^ | 1.08e^-01^ | 0.24 | 8.09e^-01^ | 8324 | 0.00 | 0.42 | **1.12e^-01^** | 0.97 | **0.005** |
| Flavonoids | EC | -5.58e^-02^ | 4.60e^-03^ | -12.1 | 8.53e^-34^ | -1073 | **0.05** | 0.59 | **1.10e^-10^** | 0.67 | **0.040** |
| Flavonoids | Circularity | 4.02e^-02^ | 4.79e^-03^ | 8.40 | 4.60e^-17^ | -1002 | 0.02 | 0.57 | **2.96e^-12^** | 0.70 | 0.106 |
| Flavonoids | Shape index | -2.73e^-02^ | 5.14e^-03^ | -5.30 | 1.17e^-07^ | -958 | 0.01 | 0.58 | **8.87e^-13^** | 0.69 | **0.001** |
| Flavonoids | Fractal dimension | -1.40e^-02^ | 4.77e^-03^ | -2.93 | 3.36e^-03^ | -944 | 0.00 | 0.60 | **5.94e^-16^** | 0.71 | **0.005** |
| Flavonoids | Area | -1.37e^-02^ | 4.78e^-03^ | -2.87 | 4.12e^-03^ | -931 | 0.00 | 0.58 | **2.09e^-15^** | 0.70 | **0.005** |
| Flavonoids | MC | -1.40e^-02^ | 4.87e^-03^ | -2.85 | 4.31e^-03^ | -940 | 0.00 | 0.59 | **3.06e^-15^** | 0.70 | **0.018** |
| NBI | Area | 1.47e^-01^ | 1.40e^-02^ | 10.5 | 1.14e^-25^ | -2179 | **0.08** | 0.72 | **2.15e^-09^** | 0.43 | 0.661 |
| NBI | EC | 1.00e^-01^ | 1.42e^-02^ | 7.07 | 1.50e^-12^ | -2122 | 0.04 | 0.68 | **1.97e^-08^** | 0.41 | 0.378 |
| NBI | Circularity | -6.32e^-02^ | 1.45e^-02^ | -4.36 | 1.28e^-05^ | -2092 | 0.02 | 0.68 | **5.03e^-07^** | 0.43 | 0.239 |
| NBI | MC | 4.01e^-02^ | 1.43e^-02^ | 2.80 | 5.11e^-03^ | -2081 | 0.01 | 0.69 | **1.54e^-08^** | 0.39 | 0.239 |
| NBI | Shape index | 4.06e^-02^ | 1.54e^-02^ | 2.63 | 8.46e^-03^ | -2080 | 0.01 | 0.68 | **1.38e^-11^** | 0.53 | 0.769 |
| NBI | Fractal dimension | 3.92e^-03^ | 1.41e^-02^ | 0.28 | 7.82e^-01^ | -2073 | 0.00 | 0.70 | **7.98e^-10^** | 0.46 | 0.466 |

Note: EC = Edge complexity, MC = Macro-shape complexity, NBI = Nitrogen balance index, SEM = Standard error of the mean, R^2^m/c = marginal/conditional explained variance (i.e., excluding/including a random effect for tree identity)

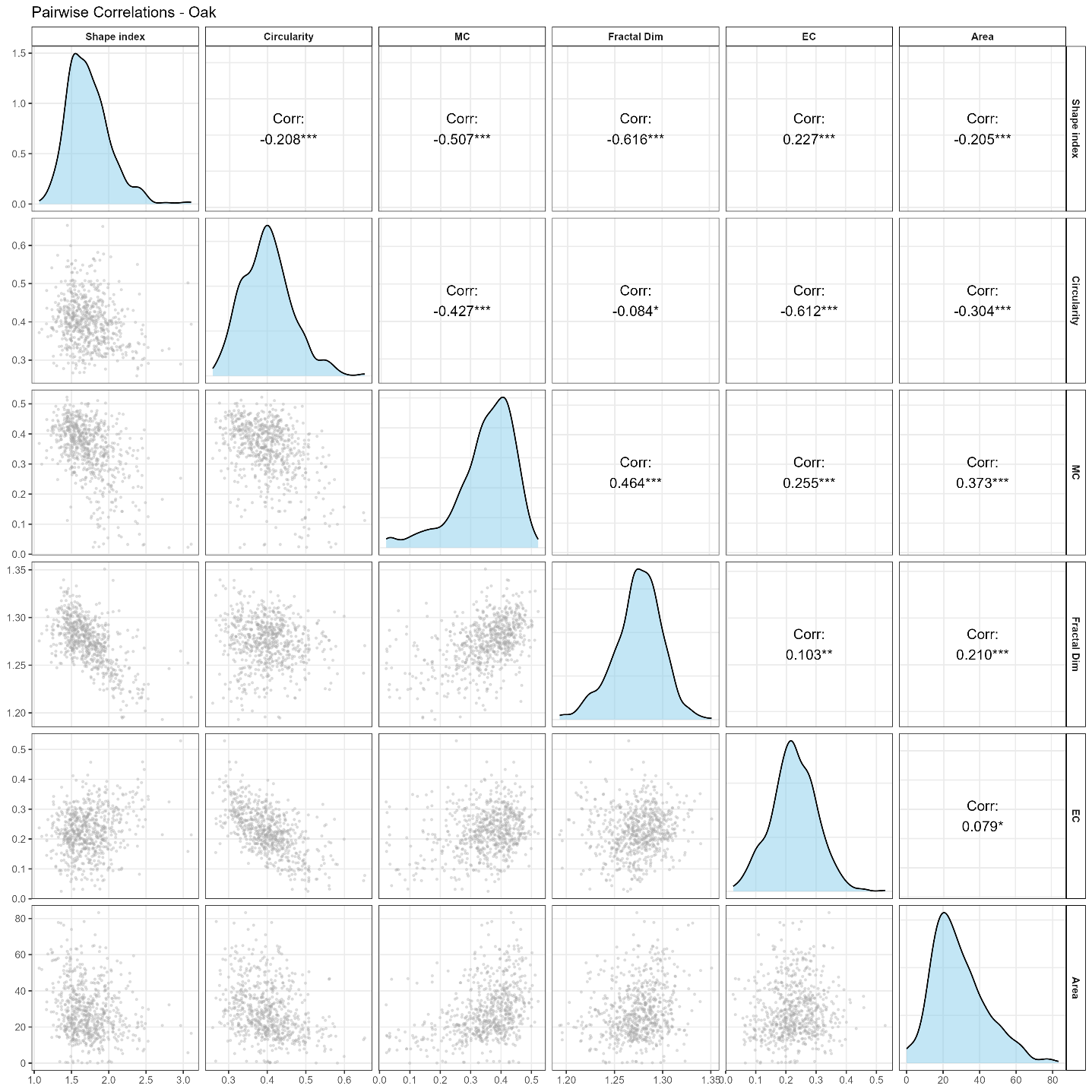

**FIGURE S1:** Correlations between all morphological metrics used in the principal component analysis (Fig. 2) from the Oak dataset.

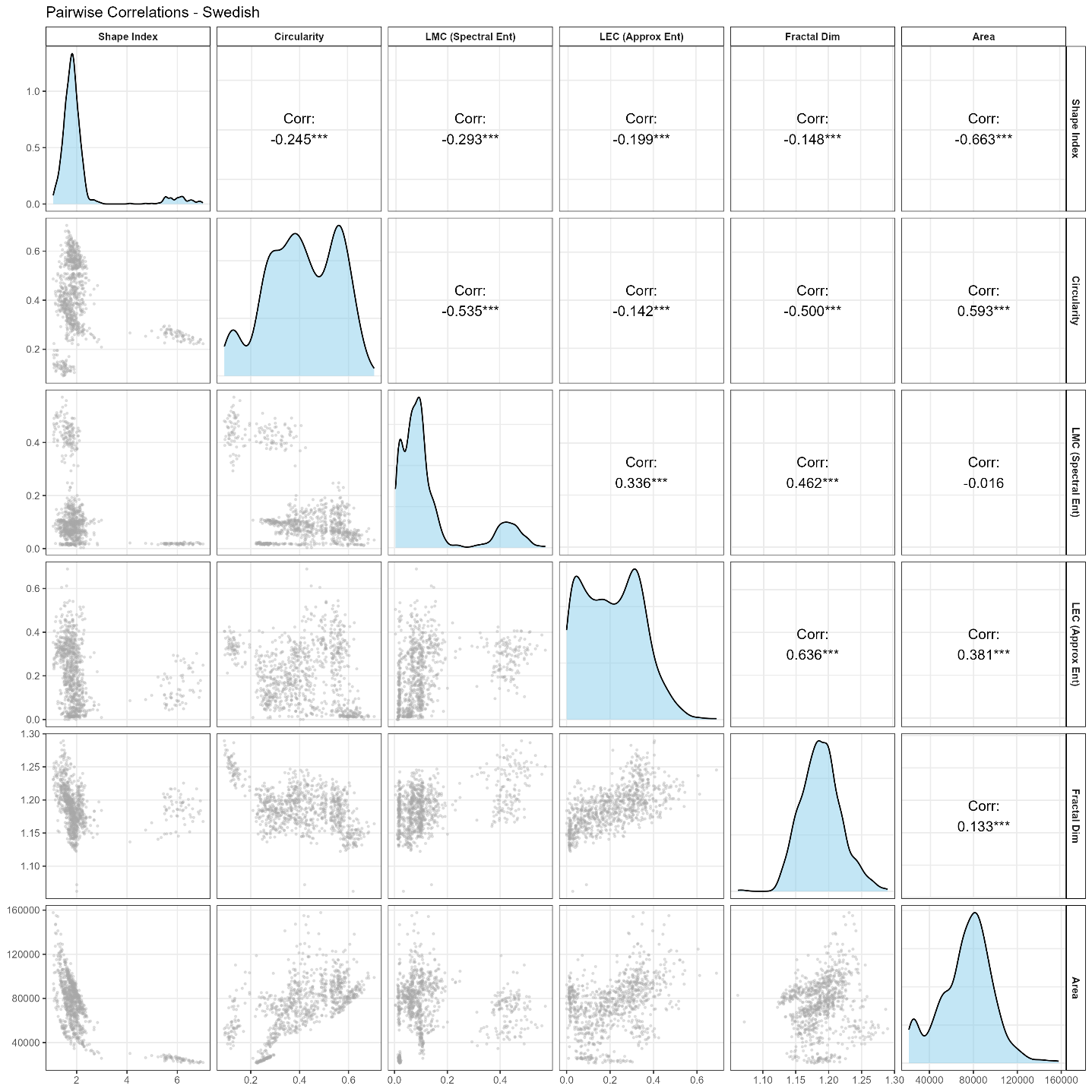

**FIGURE S2:** Correlation between all morphological traits used in the principal component analysis (Fig. 4) from the Swedish Leaf dataset.
